# Evolutionary simulations reveal role for genomic recombination in the evolution of gene regulatory network complexity and robustness

**DOI:** 10.1101/2025.08.28.672878

**Authors:** Madison Chapel, Carl G de Boer

## Abstract

The gene regulatory networks (GRNs) of eukaryotes are dramatically more complex than the GRNs of prokaryotes, but we lack a complete picture of the selective pressures that have shaped this difference. Here, we use a biochemically informed model of gene regulation to simulate GRN evolution and explore the role that reproductive strategy plays in shaping regulatory complexity. We find that recombining and non-recombining populations converge to the same level of complexity, even in the absence of selection. However, recombination modifies the rate at which complexity emerges, accelerating convergence to the complexity plateau in changing environments while slowing the process in static environments. Our results suggest that, rather than being under direct selection, regulatory complexity may emerge as a byproduct of other evolutionary processes. These results highlight how reproductive strategy and environmental change interact to influence evolutionary trajectories.

## Introduction

While many of the mechanisms underlying gene expression are similar in eukaryotes and prokaryotes, the topographies of their gene regulatory networks (GRNs) are fundamentally different^1^. Prokaryotes have TFs that are specific enough to bind only a single site in the genome^2^, and the vast majority of genes are regulated by few transcription factors (TFs), resulting in simple GRNs with few connections between TFs and genes. Meanwhile, the vast majority of eukaryotic TFs are much less specific than prokaryotic TFs. Regulation depends on combinatorial binding by multiple TFs to achieve specificity^2^, resulting in much more complex GRNs with many connections between TFs and genes^3^.

There are a number of factors that may contribute to the differences in prokaryotic and eukaryotic GRN complexity. One possibility is that these differences are not adaptive, but arose through genetic drift associated with reduced effective population sizes in eukaryotic organisms, as has been proposed for other forms of genomic complexity^4,5^. Supporting this idea, simulations have shown that the gain and loss of TF binding sites is a slow process, particularly for longer motifs, unless selection is strong (as it is in large prokaryotic populations), which could prevent eukaryotes from evolving specific motifs (which are necessarily long)^6^. However, adaptive explanations have also been proposed. For example, simulations have revealed that weak cooperative interactions between gene products tend to evolve when organisms have to perform a larger number of biological tasks^7^. As even the simplest eukaryotes have more complex cellular architectures than prokaryotes^8^, complex regulatory networks may have evolved to enable the gene expression profiles necessary to complete these tasks. Finally, eukaryotes and prokaryotes differ in the physical organization of DNA within the cell. In eukaryotes, dense chromatin restricts the number of genomic regions accessible to TFs^9^, although this restriction is not sufficient to enable the same degree of specificity as in prokaryotes^10,11^.

One possible explanation for differences in GRN complexity that has been underexplored is how differing reproductive strategies between eukaryotes and prokaryotes may influence the complexity of their regulatory networks. Prokaryotes typically reproduce asexually, resulting in clonal lineages, with mutation as the main driver of genetic variation in the population. While some additional variation can be obtained through horizontal gene transfer, most offspring are nearly identical to their parents. Eukaryotes favour sexual reproduction, with each offspring inheriting a mixture of genes from both parents through recombination. Under asexual reproduction, a tight coupling between TFs and their binding sites could evolve. In contrast, such specificity may be strongly disfavoured by a sexual reproductive strategy, as recombination will inevitably combine incompatible TF and binding site alleles. For example, a TF allele from one parent combines with a binding site allele from the other, resulting in dysregulation of the gene in the offspring. In a simple GRN, a differentially acting TF allele would change the expression of its target genes far more than in a complex GRN, where the TF is just one among many contributing to each gene’s expression **(Figure 1A)**.

**Figure 1.**
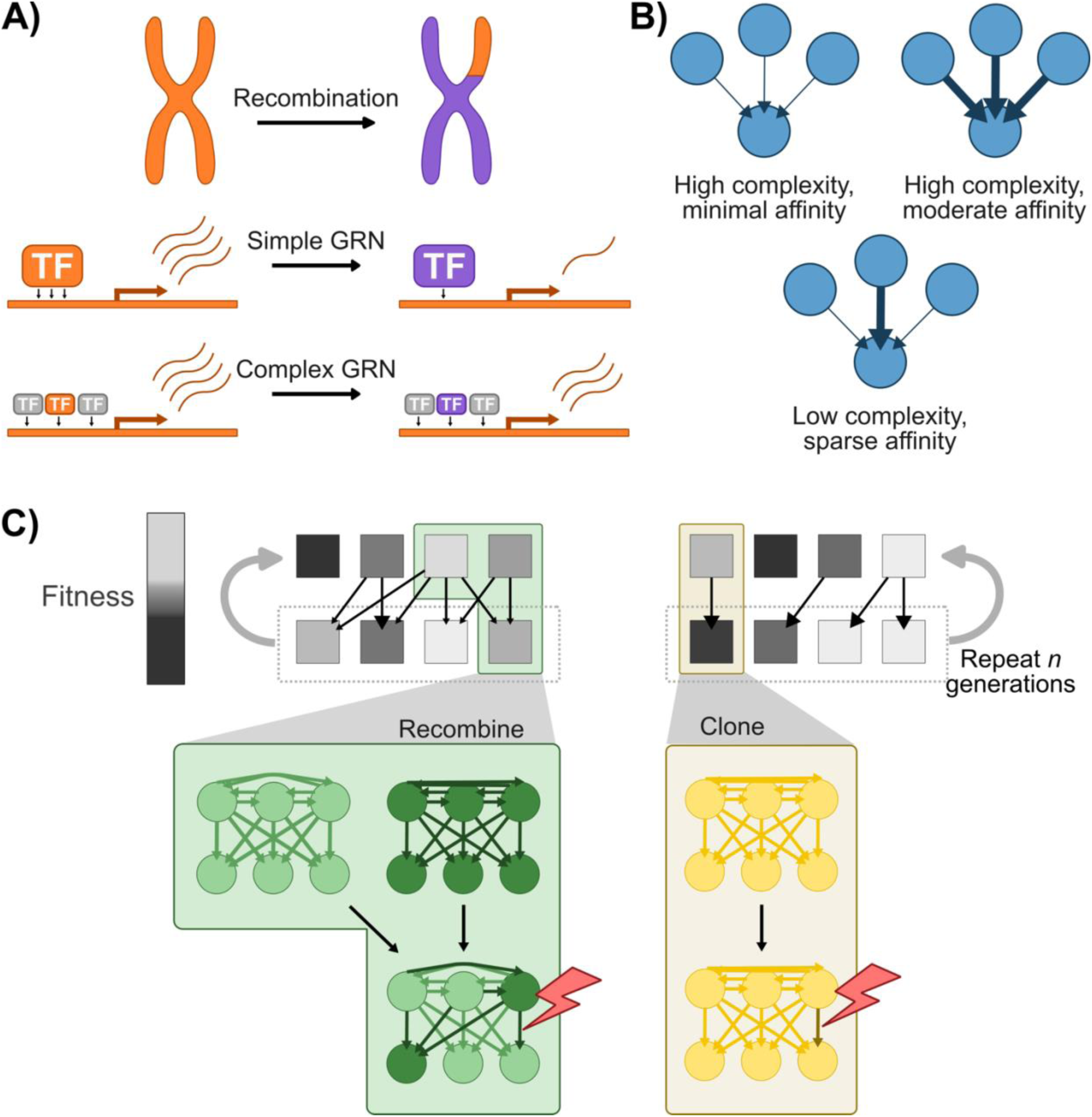
GRN model and simulation set-up. **A)** Recombination may introduce alternate TF alleles (i.e., alleles from the other parent). In a simple GRN, a differentially acting TF could alter target gene expression, while in a complex GRN, expression levels would be buffered by the activity of additional TFs. **B)** GRNs are initialized in one of three states: a high complexity state where each TF binds with minimal affinity (thin arrows), a high complexity state where each TF binds with moderate affinity (heavy arrows), and a low complexity state where a single TF binds with strong affinity while the remainder bind with weak affinity. **C)** During each generation of the simulation, individuals (squares) are selected with probability proportional to fitness (greyscale intensity). In populations with recombination (green), offspring are a hybrid of two parents, inheriting binding affinities (coloured arrows) from each. In populations without recombination (yellow), offspring are clones of parents. Offspring in both populations are subject to mutation (lightning bolt) before the process repeats.

Previous work has explored the relationship between recombination and robustness, the ability to maintain a stable phenotype after perturbations such as mutation or recombination itself^12^. Across numerous studies, recombining populations achieve higher robustness than non-recombining ones^13–19^. The underlying mechanisms that drive this increased robustness are yet to be fully understood. Singhal et al. suggested two paths by which robustness may be achieved: increased physical linkage, so that recombination doesn’t disrupt regulatory interactions, or reduced epistasis, so that loci act more independently^13^. How recombination and regulatory complexity interact to influence robustness remains an outstanding question. We hypothesized that if complexity makes populations more robust to changes introduced during recombination, recombining populations should evolve higher complexity than non-recombining populations, and more complex GRNs should exhibit greater robustness than less complex ones.

Here, we develop an *in silico* GRN model and perform evolutionary simulations to explore differences in regulatory complexity and robustness resulting from recombination. We demonstrate that recombining and non-recombining populations converge to similar complexity levels under a variety of conditions, including mutation rates, initial binding affinity strengths, and starting GRN topologies. However, recombining populations consistently demonstrate greater mutational robustness, indicating that robustness can evolve independently of GRN complexity. Finally, we reveal that, although all populations converge to the same end state of complexity, factors such as recombination and environmental changes influence the rate at which complexity emerges in populations.

## Results

### Using a biochemically informed GRN model to simulate evolution

To explore how recombination influences GRN evolution, we developed a simulation framework (**Methods**). In this simulation, there are two gene types: target genes, the expression of which determines fitness, and TFs, which regulate the target genes and each other, but do not directly impact fitness. The 200 target genes and 20 TFs are randomly arranged on a single linear “chromosome”, and the binding affinities between them are captured in a 220 x 20 matrix. A gene’s expression is the sum of TF binding probabilities (determined by the combined effect of their affinity and expression), each weighted by their effect on expression (+1 for activators, −1 for repressors; *n*=10 of each). Each target gene has a randomly initialized ‘optimal’ expression; its fitness is normally distributed with respect to the log(expression), centred at this optimum. An individual’s fitness is the product of the fitness measures for all target genes, simulating a system where all target genes are equal and essential.

In each generation, individuals are selected with probability proportional to their fitness and allowed to reproduce. Without recombination, offspring are a clonal replicate of a parent genome. With recombination, offspring are a hybrid of both parents, with a single randomly selected recombination site determining which genotypes are passed on. The TF affinities for each gene are inherited as a unit from a single parent, simulating inheritance of a single cis-regulatory region for each gene. For both with and without recombination, the TF affinities of offspring are subject to mutation. A new generation then begins, and the process repeats (**Figure 1C**). Each simulation was done in 10 replicates with a population size of 1,000 across one million generations.

### Regulatory complexity converges to similar plateaus across conditions

Using our GRN simulation framework, we first asked how varying the mutation rate impacted GRN evolution. Higher mutation rates would lead to more genetic diversity between individuals, intensifying the selective pressure for greater recombinational robustness and potentially favouring the evolution of more complex GRNs. Higher mutation rates should also increase the mutational burden at each generation, resulting in different fitness trajectories. We tested both high (∼4 per individual per generation) and low (∼1 per individual per generation) mutation rates. Populations with higher mutation rates consistently plateaued at a lower fitness due to the greater mutational burden (**Figure 2**), as expected^20^. Further, populations reproducing with recombination consistently reached higher fitness levels and evolved more rapidly than those reproducing without recombination, consistent with their increased ability to eliminate deleterious alleles^21–24^.

**Figure 2.**
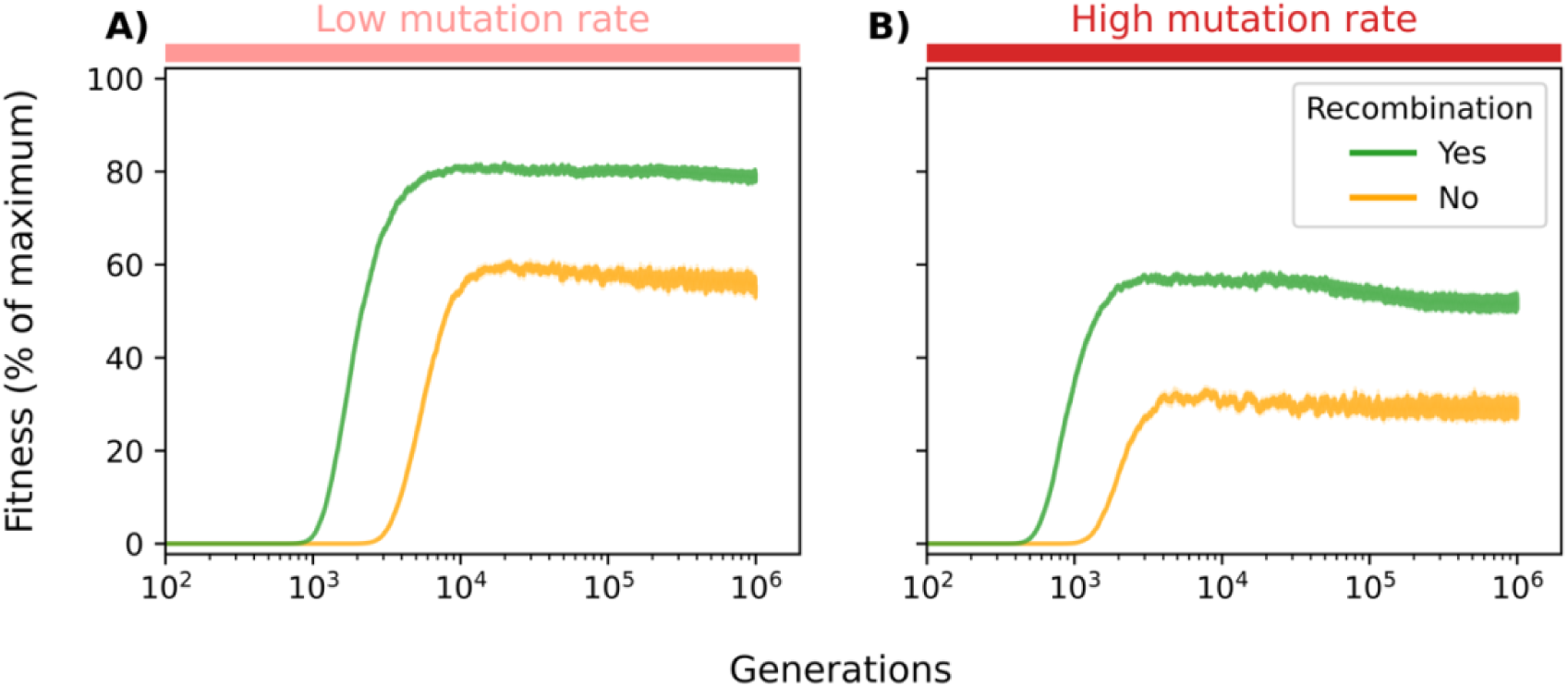
Mutation rate and recombination alter mutation-selection balance. Fitness (y-axis) shown as mean ± SEM (n = 10 replicates) for 1 million generations (x-axis). Recombining (blue) and non-recombining (yellow) populations were evolved with **(A)** low or **(B)** high mutation rates. All populations were initialized with minimal binding affinity.

We next tested how GRN initialization strategy influenced GRN evolution (**Methods**). Although previous work^14^ initialized networks with random affinities, this approach may produce an initial population with substantial fitness variability. Consequently, early generations are dominated by the fittest individuals, resulting in a large ‘founder effect’ and increased variability in simulation outcomes. To avoid this, we initialized GRNs in three different states: two high complexity states, one with minimal initial binding affinities and one with moderate initial binding affinities for all TF-TF and TF-Target pairs, and a low complexity state where expression of each TF or Target is regulated by a single randomly-selected TF with maximal binding affinity while the remaining TFs have minimal affinity (**Figure 1B, Supplementary Figure S1**). For sparse affinity initialization, the same initial GRN states were used to seed both recombining and non-recombining populations.

Despite converging to similar complexity levels (more on this below), different GRN initialization strategies led to distinct evolutionary trajectories. Here, complexity for each gene is quantified as 1 – the Gini index of TF binding probabilities^3^, with binding probabilities determined by binding affinity and TF concentration^25^. GRN complexity is then calculated as the mean complexity across all genes (**Methods**). Populations initialized in the low complexity state, where each gene is strongly bound by only a single TF, initially maintained low complexity before gradually increasing as additional regulatory interactions accumulate **(Figure 3A)**. A similar trend was observed in populations initialized with minimal affinity GRNs **(Figure 3B)**. Although these began with maximal complexity due to equal input from all TFs, a low complexity state rapidly emerges as strong binding affinities become fixed in the population. From here, complexity increases similarly to the populations initialized in the low complexity state.

**Figure 3.**
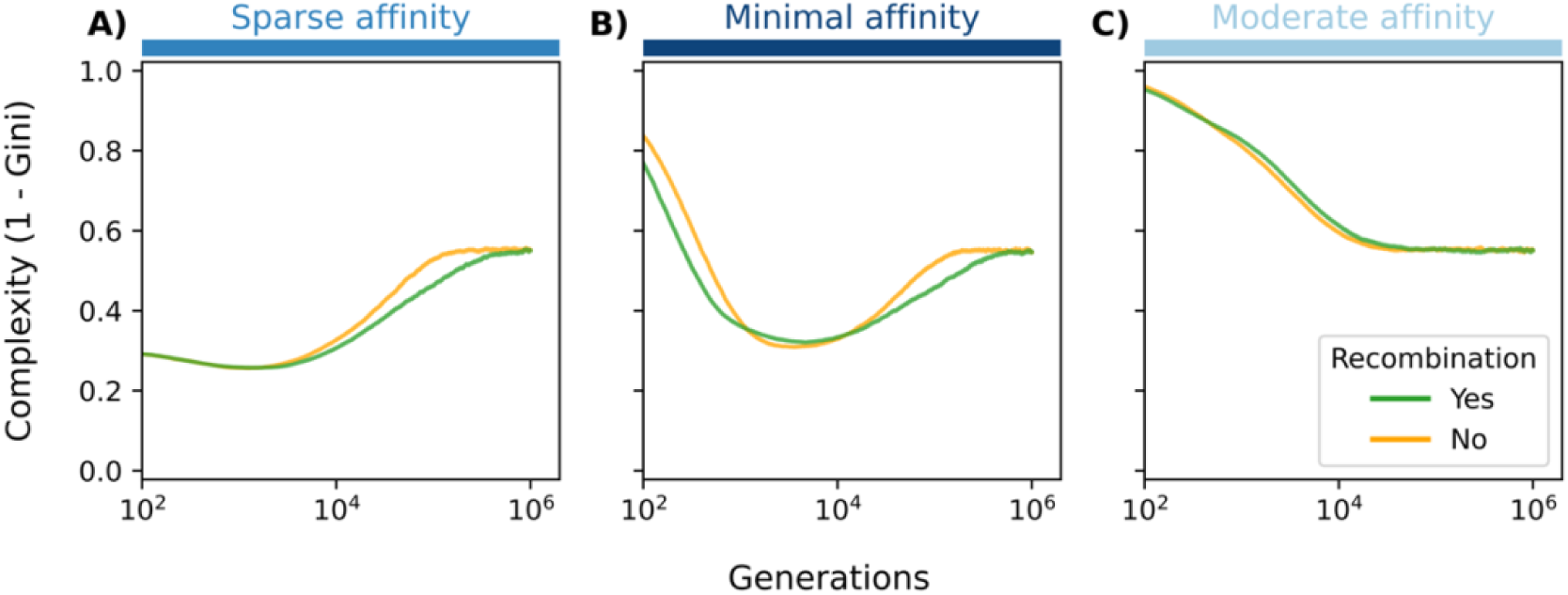
Initial binding affinities alter how complexity emerges but similar plateaus are reached. Complexity (y-axis) shown as mean ± SEM (n = 10 replicates) for 1 million generations (x-axis). Recombining (blue) and non-recombining (yellow) populations were initialized with **(A)** sparse binding affinity, **(B)** minimal binding affinity, or **C)** moderate initial affinity. All populations were evolved with high mutation rates.

In contrast, the populations initialized with moderate binding affinities followed a slightly different trajectory. Here, all activating and repressing TFs contribute moderately and equally to expression at the beginning of the simulation. Adaptive mutations are partially buffered by other regulating TFs, resulting in a gradual reduction in complexity as some edges are strengthened and others are pruned (**Figure 3C**). This buffering may reduce the individual impact of mutations, resulting in smaller fitness differences between individuals, and causing populations to reach the fitness plateau more slowly than populations initialized with minimal or sparse affinities (**Supplementary Figure S3**. Interestingly, while GRN complexity converged to a similar plateau regardless of GRN initialization, convergence occurred more rapidly in populations without recombination (**Figure 3, Supplementary Figure S4)**.

### Changing environments increase the rate at which complexity plateaus

In our previous simulations, expression goals remained constant, allowing populations to optimize their regulatory networks under stable conditions. However, selective pressures are rarely static – environments change over time, populations migrate, and other organisms create competition. Any of these scenarios could change expression optima. We reasoned that a changing environment might result in increased regulatory complexity due to the presence of lingering regulatory interactions from past adaptations. The new expression optimum could be achieved by novel regulatory interactions that counteract existing ones.

Additionally, complex regulatory networks may facilitate faster adaptation to shifting conditions through polygenic adaptation^26,27^. In a simple GRN, where a target gene is regulated by a single TF, matching a new optimum typically requires the fixation of a large-effect mutation, such as one altering the binding of a TF or one that alters expression of a regulating TF. In contrast, when a gene is influenced by multiple TFs, with each contributing relatively little, selection can act on many loci simultaneously and adaptation can proceed through small changes throughout the network that combine to have greater effects. Recombining populations are particularly well-suited to this strategy, as they can combine independently beneficial variants that may already exist as standing variation within the population^21–24^. Accordingly, we next tested whether environmental shifts could promote the evolution of regulatory complexity.

We altered our simulation by introducing environmental shifts after populations have adapted to their initial environment (150,000 generations for high mutation rate populations, 300,000 generations for low mutation rate). From then on, the expression optima change for a subset of 20 randomly selected target genes every time the population approaches the fitness plateau (**Supplementary Figure S5**). As expected, recombining populations adapted to new environments faster than non-recombining ones, requiring an average of 1,112 and 2,289 generations, respectively, to regain fitness after each shift (**Supplementary Figure S6**). Despite repeated environmental changes, regulatory complexity ultimately plateaued at a similar level across all conditions, stabilizing at around ∼0.55 – the same value observed previously in the static environments. However, for populations that had not yet reached this level when environmental shifts were introduced, the changing environment accelerated the rate at which complexity increased. This was particularly pronounced in recombining populations, where a distinct inflection point can be seen when environmental changes are introduced (**Figure 4B, Supplementary Figure S7,S8**).

**Figure 4.**
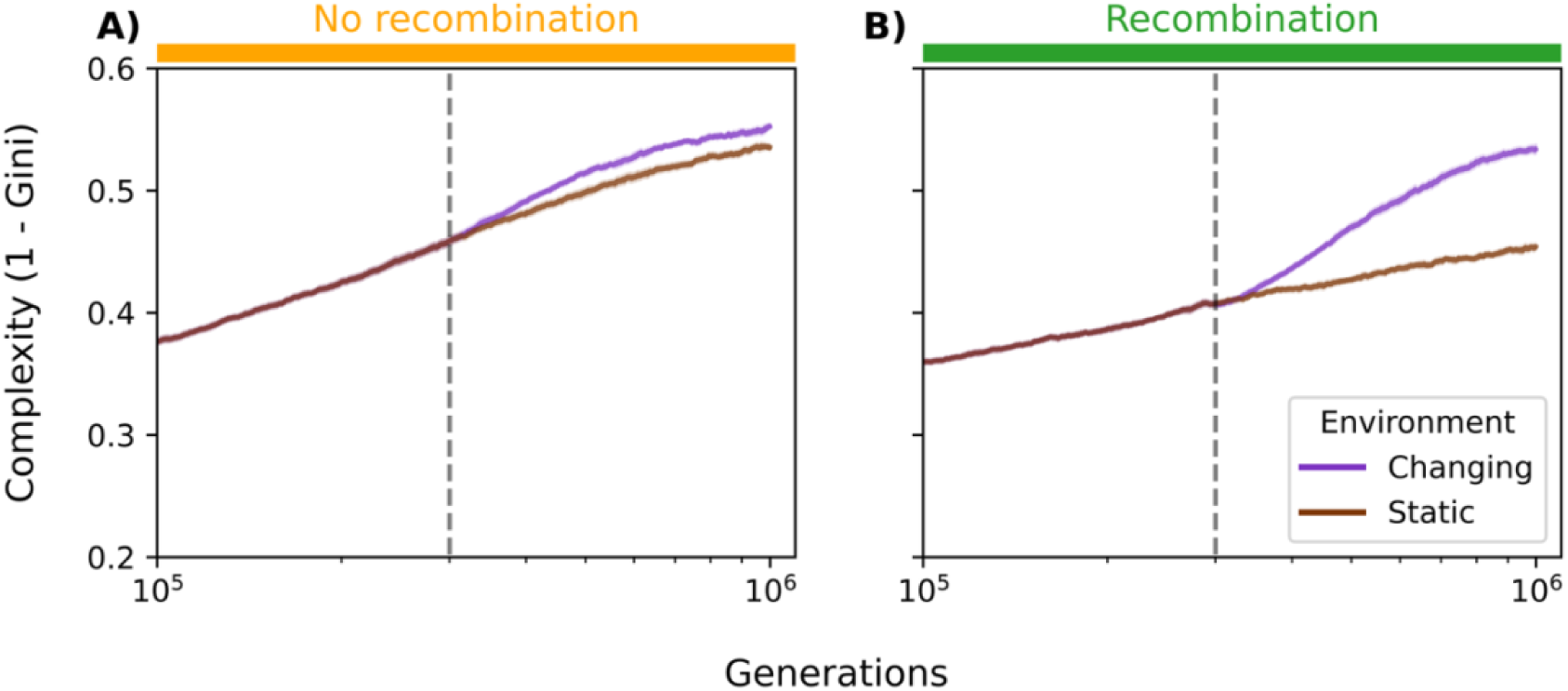
Complexity emerges more rapidly during environmental changes, especially for recombining populations. Complexity (y-axis) shown as mean ± SEM (n = 10 replicates) up to 1 million generations (x-axis) for populations under static (green) or changing (purple) environmental conditions. Populations were evolved either **(A)** without recombination or **(B)** with recombination. All populations were initialized with minimal binding affinities and evolved with low mutation rates. Environmental changes begin at the generations marked by vertical dashed lines.

### Recombining populations achieve higher robustness

We next investigated whether recombination increased robustness, and whether mutational robustness increased with increasing regulatory complexity. To test this, we perturbed individual GRNs by introducing mutations. ‘Robustness’ was measured by determining the mean Euclidean distance between the original expression profile and 100 mutated profiles (**Methods**). Despite converging at similar complexity levels, recombining populations consistently evolved greater mutational robustness than non-recombining ones (**Figure 5, Supplementary Figure S9**), as demonstrated in previous studies^14–19^. Furthermore, no correlation was found between complexity and robustness (Pearson’s *r*^2^ <0.014 for all populations tested; **Supplementary Figure S10**). The trend of greater robustness in recombining populations was observed regardless of mutation rate or initialization conditions, although these variables did affect the robustness in different ways, especially at earlier generations (**Figure 5**).

**Figure 5.**
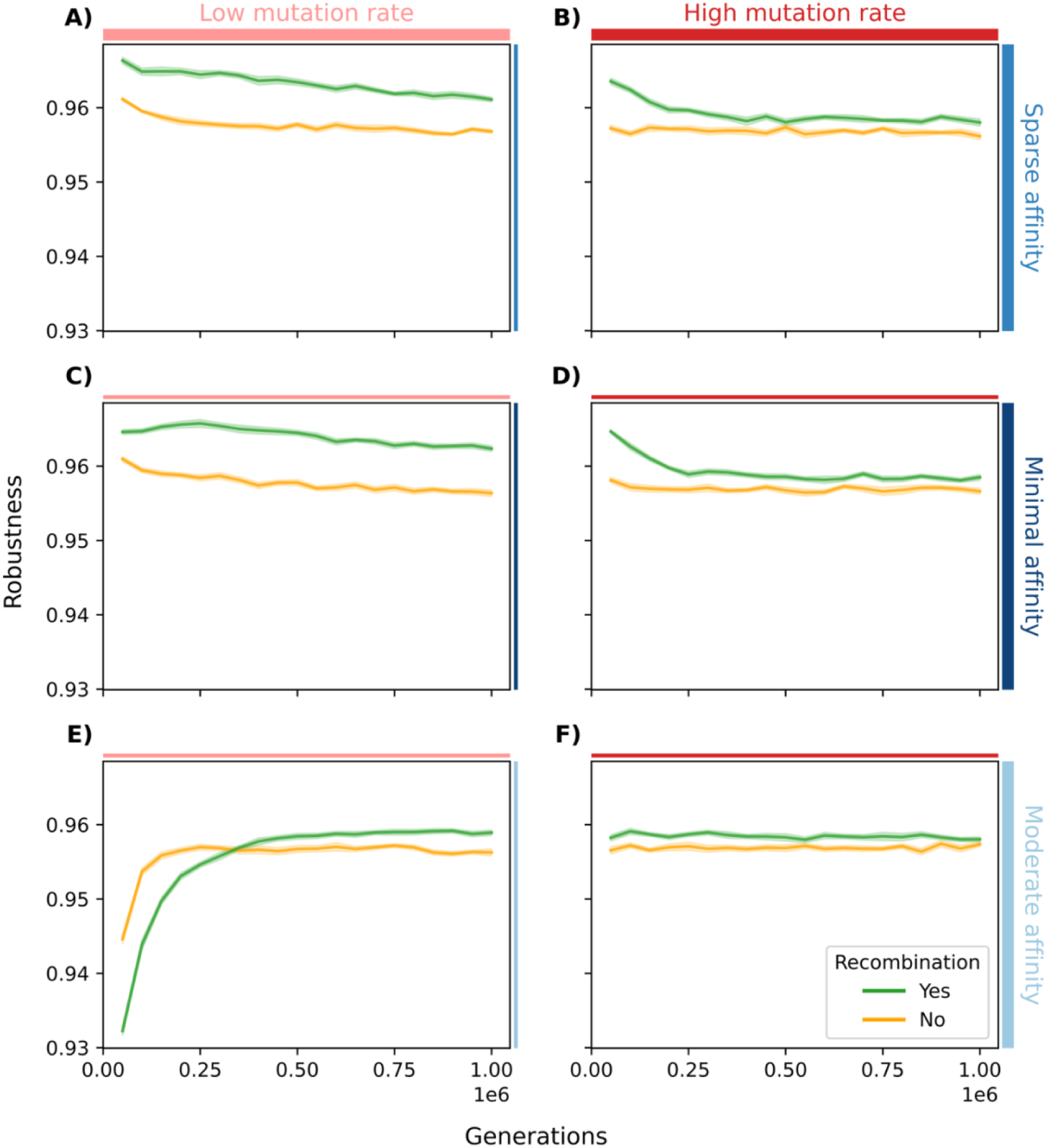
Recombining populations consistently evolve higher robustness than non-recombining. Mutational robustness (y-axis) shown as mean ± SEM (n = 10 replicates) for 1 million generations (x-axis). Populations were evolved with (blue) or without (yellow) recombination. Panels show: **(A, C, E)** Low mutation rate, **(B, D, E)** high mutation rate, **(A, B)** sparse initial affinity, **(C, D)** minimal initial affinity, **(E, F)** moderate initial affinity.

**Figure 6.**
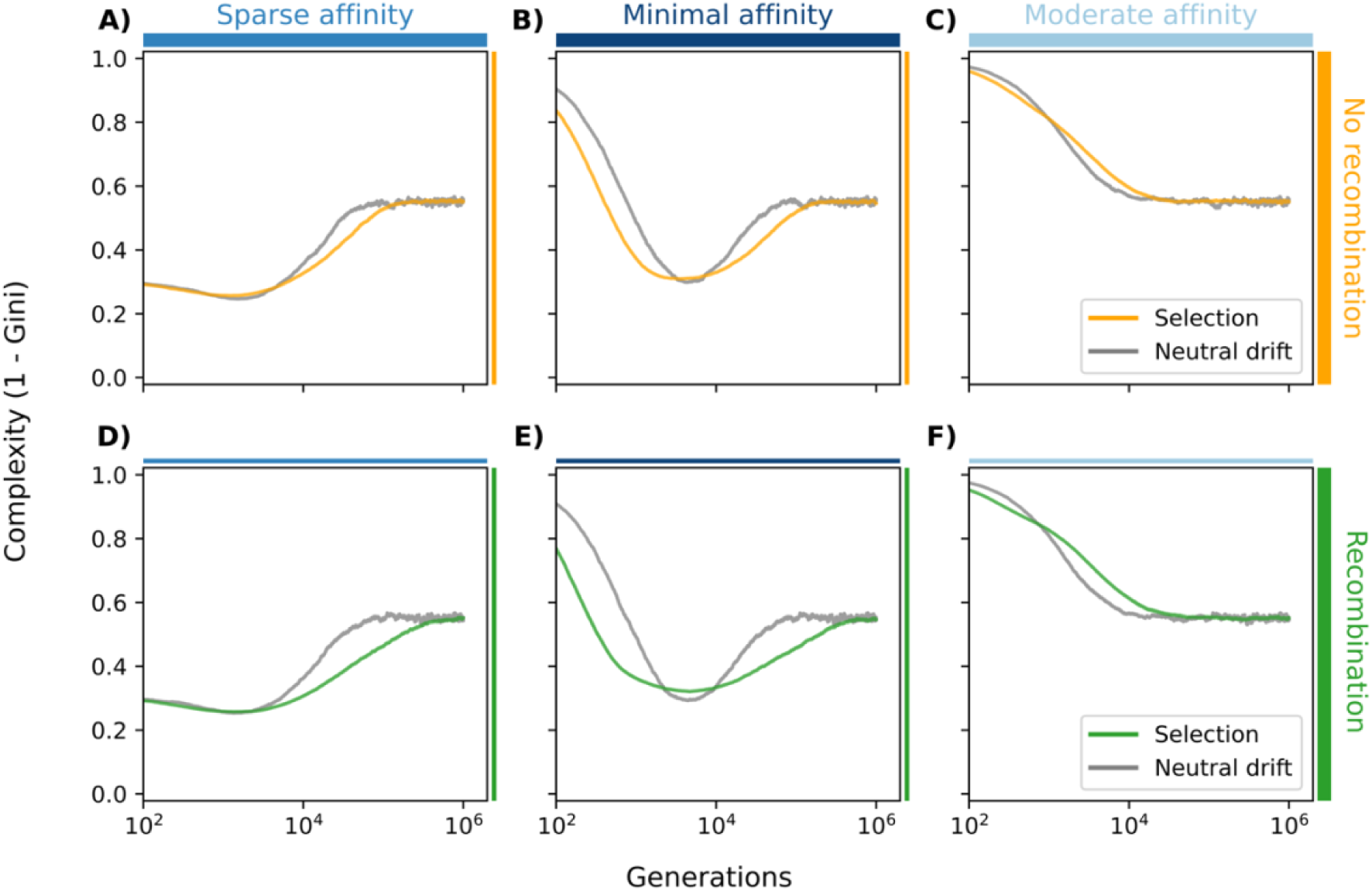
Complexity plateaus more rapidly in neutrally evolving populations. Complexity (y-axis) shown as mean ± SEM (n = 10 replicates) for 1 million generations (x-axis). Populations were evolved with (colors) or without (grey) selection. Panels show: **(A, D)** Sparse initial affinity, **(B, E)** minimal initial affinity, **(C, F)** moderate initial affinity, **(A, B, C)** no recombination, **(D, E, F)** recombination. All populations were evolved with high mutation rates.

### Similar regulatory complexity is produced by neutrally evolving populations

To determine whether the consistently observed complexity plateau represents an adaptive optimum or a neutral byproduct, we evolved populations without selection. Instead of selecting individuals with probability proportional to their fitness, individuals were selected randomly at each generation. All other parameters (i.e., mutation rate, initial binding affinities, recombination) were matched to previous simulations. These simulations represent an evolutionary null, providing an expectation of regulatory complexity in the absence of selection.

While the neutrally evolving populations converge to the same complexity as matched populations under selection, they did so more rapidly than with selection. In static environments, selection thus appears to delay this convergence, an effect that is more pronounced when recombination is present.

## Discussion

In our simulations, recombining populations consistently evolved GRNs that were more robust to mutational perturbation. This finding aligns with previous work demonstrating smaller mutational effects in recombining finite populations^13–19^. Our simulations also captured the widely-demonstrated effect of recombining populations adapting to new environments faster than non-recombining ones^28–35^.

Our simulations suggest that recombination and changing environmental pressures can influence the emergence of regulatory complexity. Reproductive mode alone did not drive the evolution of more complex GRNs, as recombining and non-recombining populations tended to plateau at similar levels of regulatory complexity. Even when selection was removed entirely, complexity still reached the same plateau. However, recombination did influence the rate at which complexity plateaued. In static environments, populations under selection plateaued more slowly than neutrally evolving populations, with this effect being most pronounced in recombining populations. When the environment was dynamic, this effect was reversed; recombination drove populations to the plateau more rapidly than in non-recombining populations.

These results suggest that selection does not act directly on complexity. Instead, complexity arises as a byproduct of how recombination and selection shape the exploration of genotype space. In static environments, when populations are near the fitness optimum, most mutations will be neutral or deleterious. Recombination more effectively purges these deleterious variants from the population, maintaining a stable phenotype but also limiting the population’s ability to navigate the space of viable regulatory networks, slowing the accumulation of regulatory complexity. In contrast, when selection pressures change, recombination promotes rapid adaptation by combining beneficial variants from otherwise unfit backgrounds^36,37^. In our simulations, complexity emerged most rapidly for recombining populations in changing environmental conditions, suggesting that when selection pressures shift, recombination instead promotes exploration of genotype space. Thus, the changes in complexity may reflect changes in how recombination mediates adaptation rather than selection for complexity.

The consistent complexity plateau across our simulations raises further questions: does this represent a biologically meaningful upper limit on regulatory complexity, or is it an artifact of the model’s constraints? Future work should aim to refine GRN models with additional regulatory mechanisms to more accurately reflect biological systems. This, along with testing a wider range of mutation rates, recombination frequencies, and other parameters, will provide further insights into how differences in regulatory complexity emerge and whether complexity itself provides an adaptive benefit. Consistent with previous simulations of gene regulatory evolution, there may be minimal selection on complexity, and instead the complexity of extant regulatory sequences may simply reflect sampling from the space of solutions that satisfies the expression constraints^3,38^. Instead, the different complexities of prokaryote and eukaryote GRNs could simply reflect the different gene regulatory hardware they utilize. While we simulated recombining and non-recombining populations with identical hardware, the reality is that prokaryotic TFs tend to be far more specific than eukaryotic TFs^2^. The question then becomes, why do prokaryotes have such specific TFs relative to eukaryotes?

While our GRN model draws on established literature to simulate regulatory networks in a biologically informed manner^3,25^, several simplifications were made to streamline the design and improve the model’s interpretability. In our model, TFs functioned exclusively as activators or repressors. In reality, TFs can have both activating and repressing activity in different contexts, can be conditionally active only after receiving an appropriate signal^39^, and these activities can change over evolutionary time. Further, the number of TF and Target genes was fixed, preventing regulatory networks from evolving via gene duplication – a process partially responsible for the redundancy in eukaryotic binding motifs^40^. Mutations in TF coding sequences, which were not simulated here, could have even more widespread effects, altering the activity of that TF throughout the genome. To make our simulations computationally tractable, we tested only a single, constant, population size of 1000 individuals. Future simulations may explore whether drift in small populations or bottlenecks of reduced population size influence the emergence of regulatory complexity.

## Methods

### Modeling GRNs

#### Binding affinity matrix

Individual GRNs were modeled as a matrix of binding affinities bounded between ln(*k*) = −5.0 and ln(*k*) = 5.0, with 20 TFs and 200 target genes. Half of the TFs were designated as activators and half as repressors. TF and target genes were randomly distributed throughout a single linear chromosome. TFs regulate the expression of all target genes and each other, resulting in a 20 x 220 matrix whose elements ln(*k_i,j_*) describe the binding affinity of the TF product of gene *i* to the promoter of gene *j*.

#### Gene expression

The binding probability *R_i,j_* of TF *i* to gene *j* was adapted from Granek and Clarke^25^:

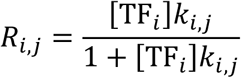

Where [TF*_i_*] is the expression level of TF*_i_* and *k_i,j_* the binding affinity of TF*_i_* to gene *j*.

The total regulatory activity *R_j_* of all TFs acting on gene *j* was calculated as the sum of the binding probabilities of all activators minus the sum of the binding probabilities for all repressors:

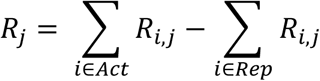

Where *Act* is the set of all activating TFs while *Rep* is the set of repressing TFs. The regulatory activity was then passed into a sigmoidal function to determine the expression level *E* in log space:

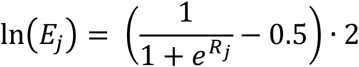

Expression was calculated iteratively until expression levels for all genes had stabilized (differences of <1 x 10^-4^ between iterations). For the first iteration, log expression levels were randomly sampled uniformly between −1 and 1, with the same initialization applied to every individual in the population.

Each of the 200 target genes was assigned an optimal expression level, 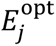. These values were drawn from a uniform distribution in log space:

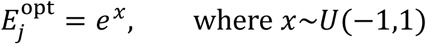

#### Fitness

The fitness of an individual, *F*, was calculated as the product of the fitness of all target genes. The fitness of each target gene was defined by how closely its expression level, *E_j_*, matched the optimal expression level, 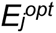, using a Gaussian function with a standard deviation of σ = 0.75. Expression levels were compared in log space:

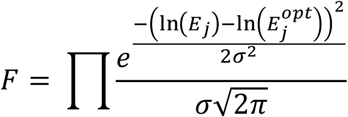

#### Complexity

Complexity of a GRN was measured using the Gini coefficient^3^. While typically used in economics to quantify inequalities in wealth distribution, the Gini coefficient can also model other inequalities within a population, such as the degree of inequality in regulatory interactions of TFs in a regulatory network. A Gini coefficient of 0 represents a distribution where all members of the population have equal wealth, while a coefficient of 1 represents a distribution where all wealth is controlled by a single individual. Complexity of regulatory interactions is measured as 1 – Gini, and ranges from 0 (expression is controlled by a single TF) to 1 (all TFs contribute equally to expression). Complexity, *C*, for each gene was measured using the binding probabilities, *R*, sorted in ascending order, of *n* TFs acting on it:

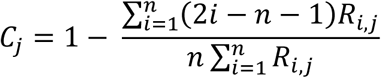

Complexity for each individual was measured as the mean complexity of all genes.

#### Robustness

Robustness was measured by introducing mutations to a GRN and measuring changes in expression. Mutation effect sizes were sampled from a Gaussian distribution with standard deviation of 2.0; the same range used for mutations during evolution simulations (see below). The change in expression was measured as the Euclidean distance between the vector of initial expression pattern and the expression pattern after mutation, compared in log space. Robustness, ρ, was measured every 50,000 generations across *k =* 100 mutations:

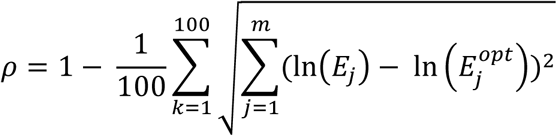

### Evolution simulations

Evolution was simulated using a genetic algorithm (GA) implemented with the DEAP Python package^41^. For all simulations, the population size was 1000 and reproduction probability was 0.5. GRNs within a population were initialized in one of three states: a high complexity state with minimal initial binding affinities for all TF-TF and TF-Target pairs, a high complexity state with moderate initial binding affinities, and a low complexity state with sparse affinities where expression of each TF or Target is regulated by a single randomly-selected TF with maximal binding affinity while the remaining TFs have minimal affinity **(Supplementary Figure S1)**.

The moderate binding affinity (*k*) was defined as ln(*k*) = 0.0. For the minimal affinity initialization, our goal was to evolve GRNs from a state without regulatory interactions. However, because affinities were represented in log space, it was necessary to determine a sufficiently low cut-off for a minimal binding affinity value. A value of 0 (no affinity) cannot be represented in log space, and choosing too low a value would result in GRNs evolving slowly or not at all (e.g., a mutation from ln(*k*) = −10.0 to ln(*k*) = −9.0 may not alter expression enough to confer a fitness advantage). The minimal initial binding affinity was set such that initial expression values varied minimally from the basal expression level of ln(*E*) = 0.0. To achieve this, we tested a range of binding affinities and selected the largest value that yielded expression levels with a standard deviation ≤ 0.025 in log space. This value depended on the number of TFs in the network, and was determined to be ln(*k*) = −(0.517 · ln (*n*) + 2.516), where *n* is the number of TFs (in our case of 20 TFs, ln(*k*) = −4.06) **(Supplementary Figure S2)**. For the high complexity initialization, a single maximal initial binding affinity was set as ln(*k*) = (0.517 · ln (*n*) + 2.516), which, with 20 TFs, is ln(*k*) = 4.06.

For populations under selection, selection was performed using a roulette function, where the probability of selection is proportional to fitness. For neutrally evolving populations, selection was performed randomly where each individual had equal probability of being selected.

For recombining populations, a custom one-point crossover function was used to independently generate two distinct recombinant offspring for each mating pair. Each offspring inherited all binding affinities prior to a randomly selected crossing-over point from one parent, and all binding affinities after the cross-over point from the other parent. The process was repeated with a new cross-over point to generate the second offspring.

Mutation effect sizes were sampled from a Gaussian distribution with standard deviation of 2.0, allowing minimal or maximal expression to be achieved within two mutations, as previously observed for yeast promoters^3^. Mutation effects were applied to the binding affinities (*k*_*i*,*j*_) in logarithmic space. The mutation probability was set such that each new individual would receive an expected 4.4 (*p* = 1.0 × 10^−3^, high mutation rate) or 1 (*p* = 2.2727 × 10^−4^, low mutation rate) mutations.

Each GA was run for 1 million generations. A total of 10 replicate populations were evolved for each set of conditions. Optimal expression goals were randomly initialized for each replicate and matched across populations with or without recombination.

To simulate selection in changing environmental conditions, simulations were resumed after fitness levels had plateaued (150,000 generations for high mutation rate simulations, 300,000 generations for low mutation rate). The mean fitness of the population was determined as a percentage of the maximum possible fitness. 20 target genes were selected at random, and their optimal expression values shifted by values sampled from a Gaussian distribution with standard deviation of 0.25, simulating a change in the environment and selection for a new expression pattern. New optimal expression values were bounded between ln(−0.9) and ln(0.9) to prevent the emergence of difficult to attain expression goals (e.g., requiring maximum expression for all target genes). When the population’s mean fitness was within a percentage point of the original mean fitness, the environmental shift was repeated.

## Acknowledgements

We thank S. Otto and J. Dennis for helpful discussions. This research was supported by the Natural Sciences and Engineering Research Council of Canada (RGPIN-2020-05425), and the Canadian Institute for Health Research (PJT-180537). M.C. was supported by a UBC IGF. C.G.D. is a Michael Smith Health Research BC Scholar.

## Data availability

Code used to run simulations and analyze results can be found at https://github.com/de-Boer-Lab/GRN-evolution/.

## Supplementary

### Figures

**Supplementary Figure S1.**
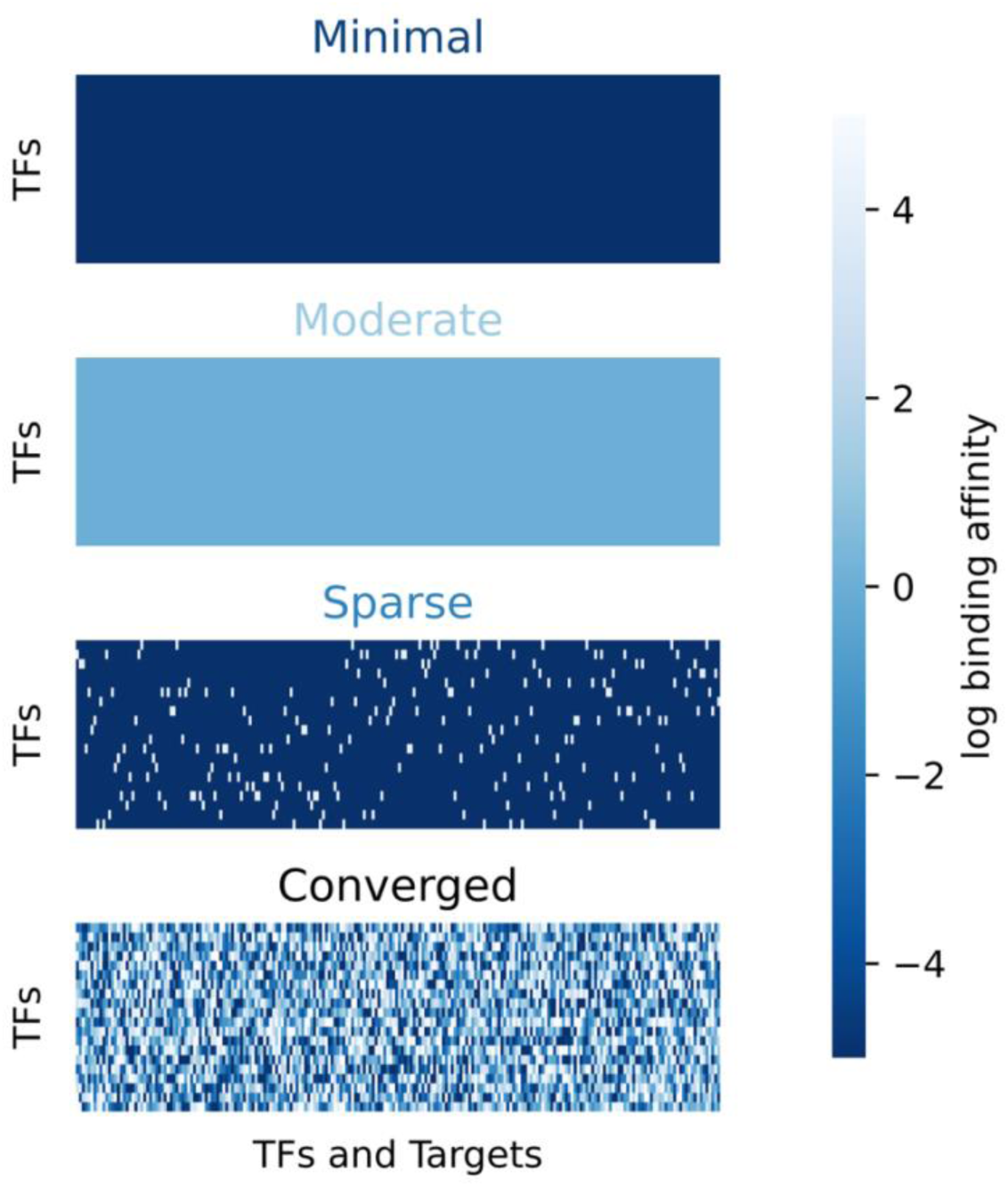
Sample GRNs in various states. GRNs are represented as a matrix of log binding affinities (colour) for 20 TFs (y-axis) regulating all other TFs and target genes (x-axis). Shown here are GRN log binding affinities (color) initialized with minimal, moderate, and sparse binding affinities, and a GRN that has converged at the complexity plateau. When initialized with sparse binding affinity, a single TF per column binds with maximal affinity while the remainder bind with minimal affinity.

**Supplementary Figure S2.**
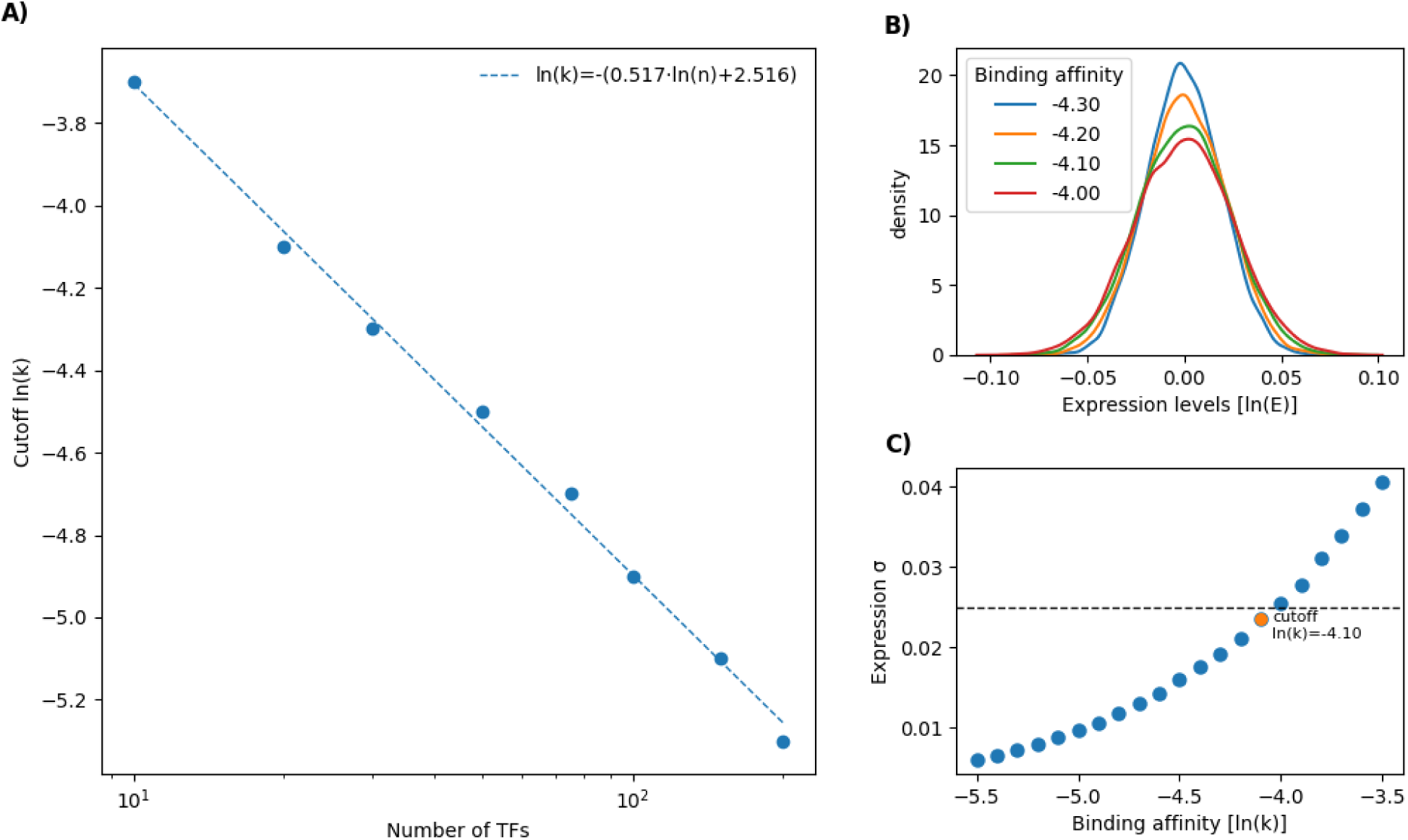
Determining minimal binding affinity initialization. **(A)** Relationship between sufficiently low binding affinity (ln(*k*)) values (y-axis) and number of TFs in a network (x-axis). A regression line was fit to cut-off values (points) for varying numbers of TFs. **(B)** Distributions of gene expression values (x-axis) for a network with 20 TFs initialized with varying binding affinities. **(C)** Relationship between binding affinity strength (x-axis) and standard deviation of expression (y-axis) for a network of 20 TFs. The threshold of σ = 0.025 is indicated with a dashed line, and the largest binding affinity value (ln(*k*) = −4.10) not exceeding the threshold is indicated in orange.

**Supplementary Figure S3.**
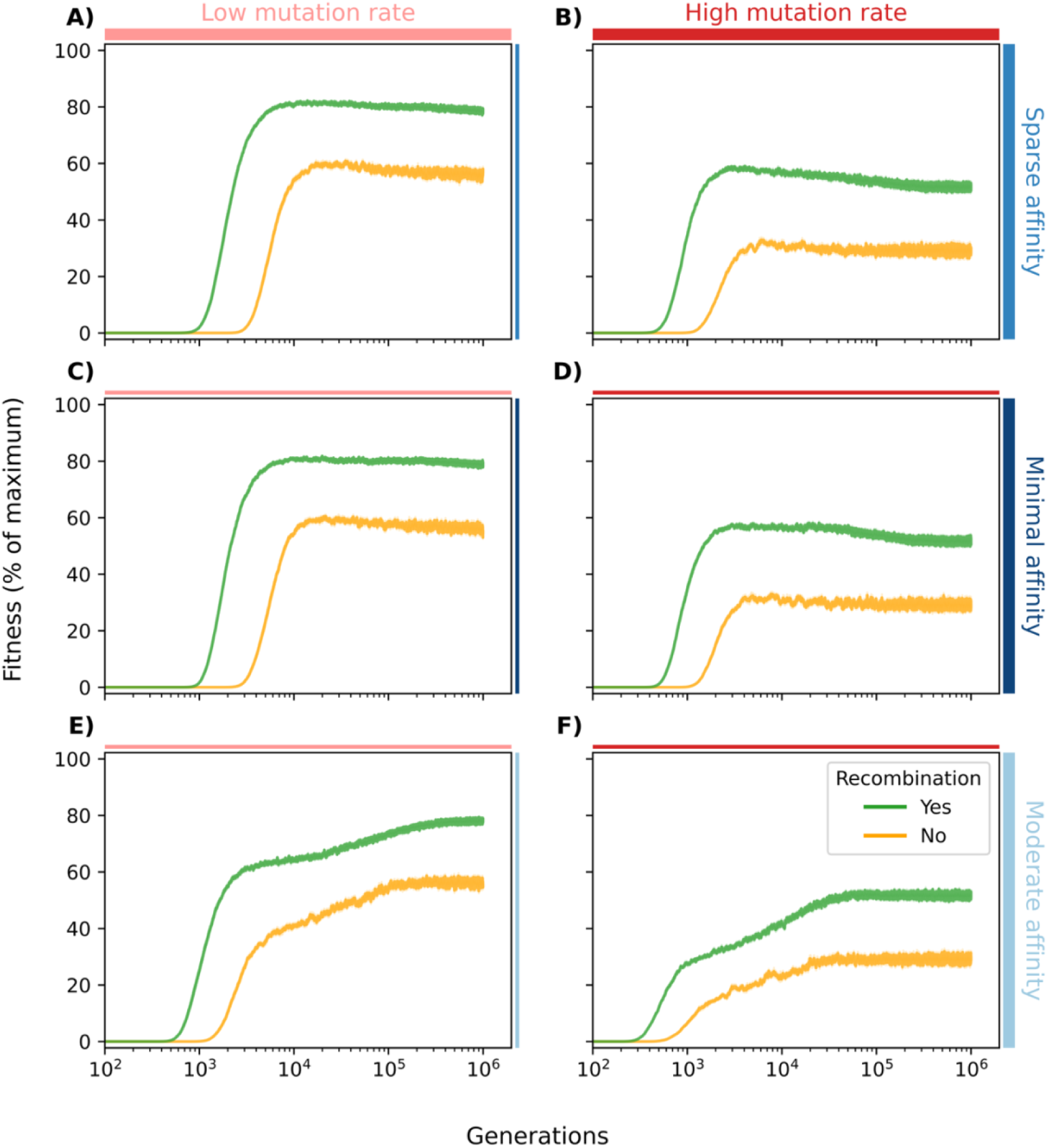
Fitness trajectories for all simulations in static environments. Fitness (y-axis) shown as mean ± SEM (n = 10 replicates) for 1 million generations (x-axis). Populations were evolved with (blue) or without (yellow) recombination. Panels show: **(A, C, E)** Low mutation rate, **(B, D, E)** high mutation rate, **(A, B)** sparse initial affinity, **(C, D)** minimal initial affinity, **(E, F)** moderate initial affinity.

**Supplementary Figure S4.**
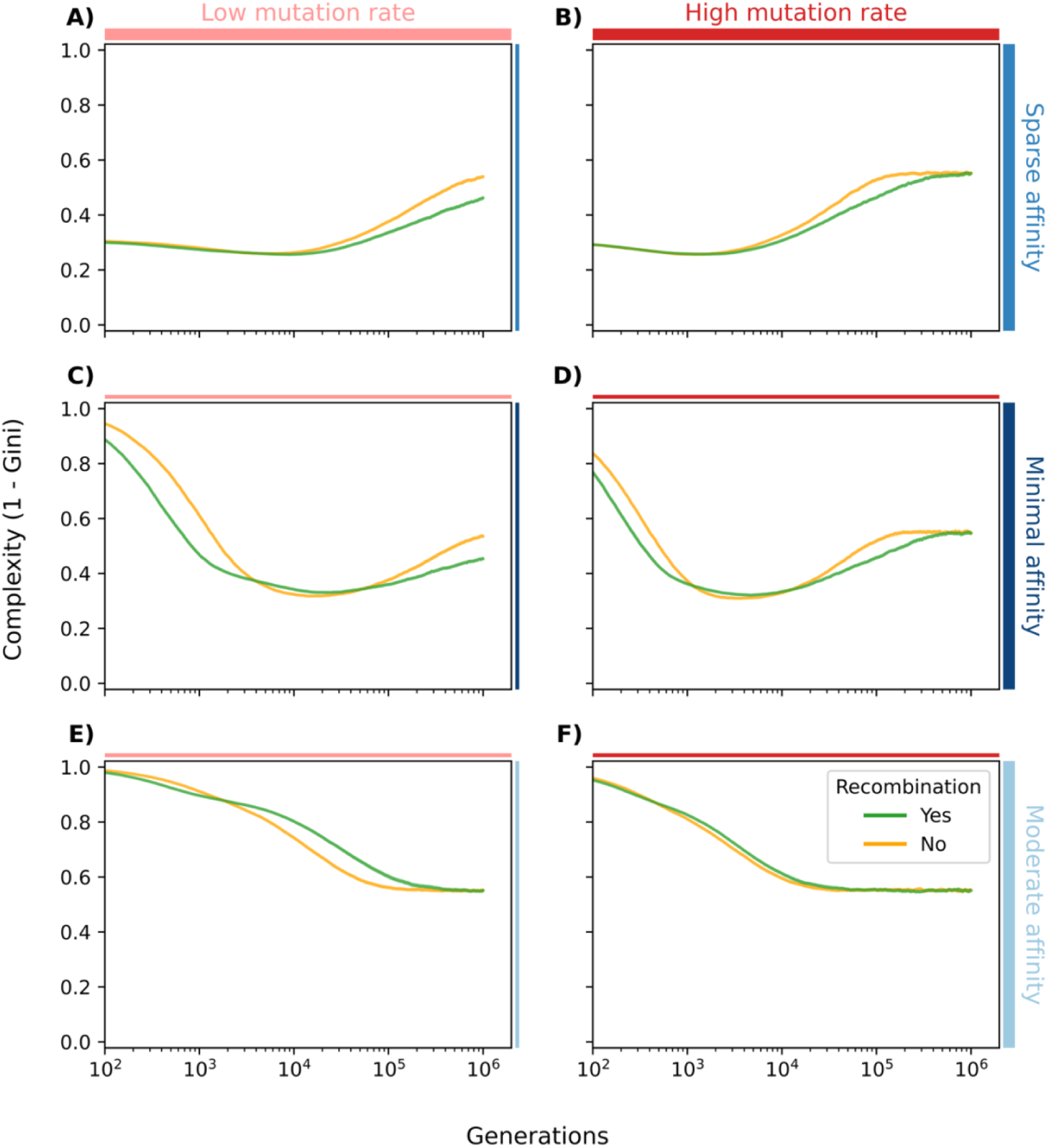
Complexity trajectories for all simulations in static environments. Complexity (y-axis) shown as mean ± SEM (n = 10 replicates) for 1 million generations (x-axis). Populations were evolved with (blue) or without (yellow) recombination. Panels show: **(A, C, E)** Low mutation rate, **(B, D, E)** high mutation rate, **(A, B)** sparse initial affinity, **(C, D)** minimal initial affinity, **(E, F)** moderate initial affinity.

**Supplementary Figure S5.**
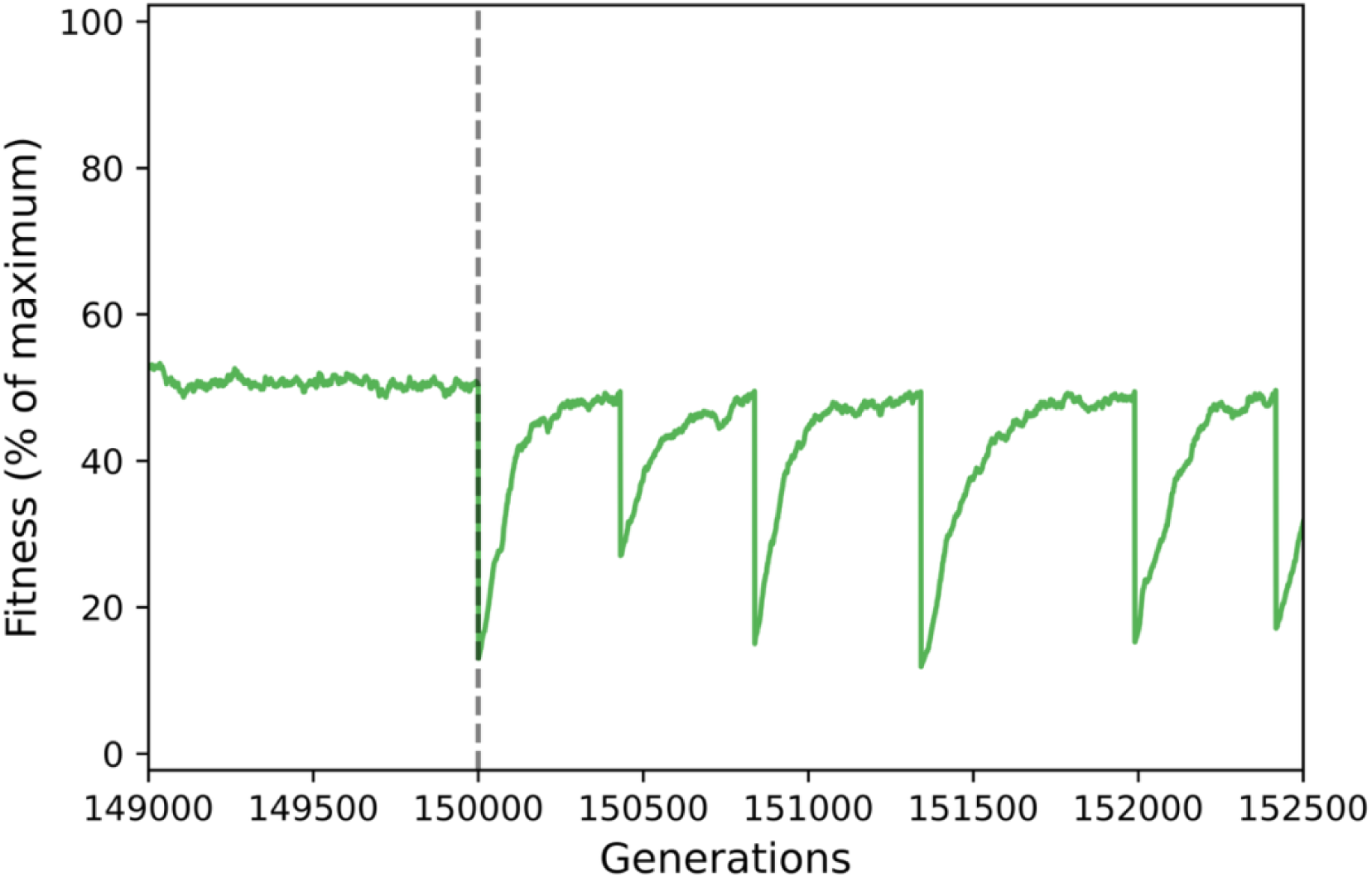
Environmental changes cause fitness to drop and recover. Example of average fitness (y-axis) evolution for a single replicate population during environmental changes, beginning at generation (x-axis) 150,000. Population was initialized with moderate binding affinity and evolved with recombination and a high mutation rate.

**Supplementary Figure S6.**
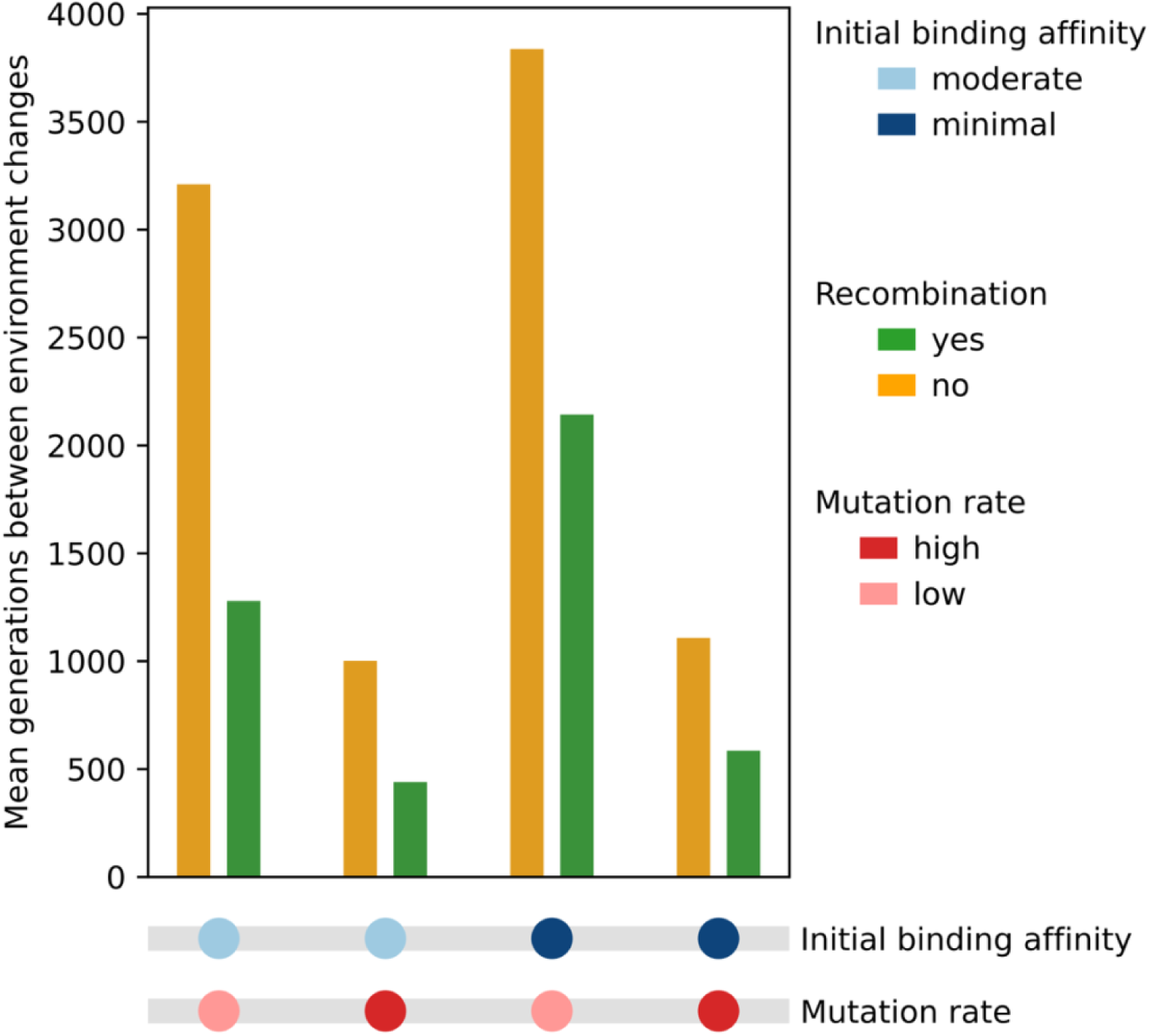
Recombining populations recover more quickly after environmental changes. Mean generations between environment changes (y-axis) for recombining (green) and non-recombining (yellow) populations across different test conditions (x-axis).

**Supplementary Figure S7.**
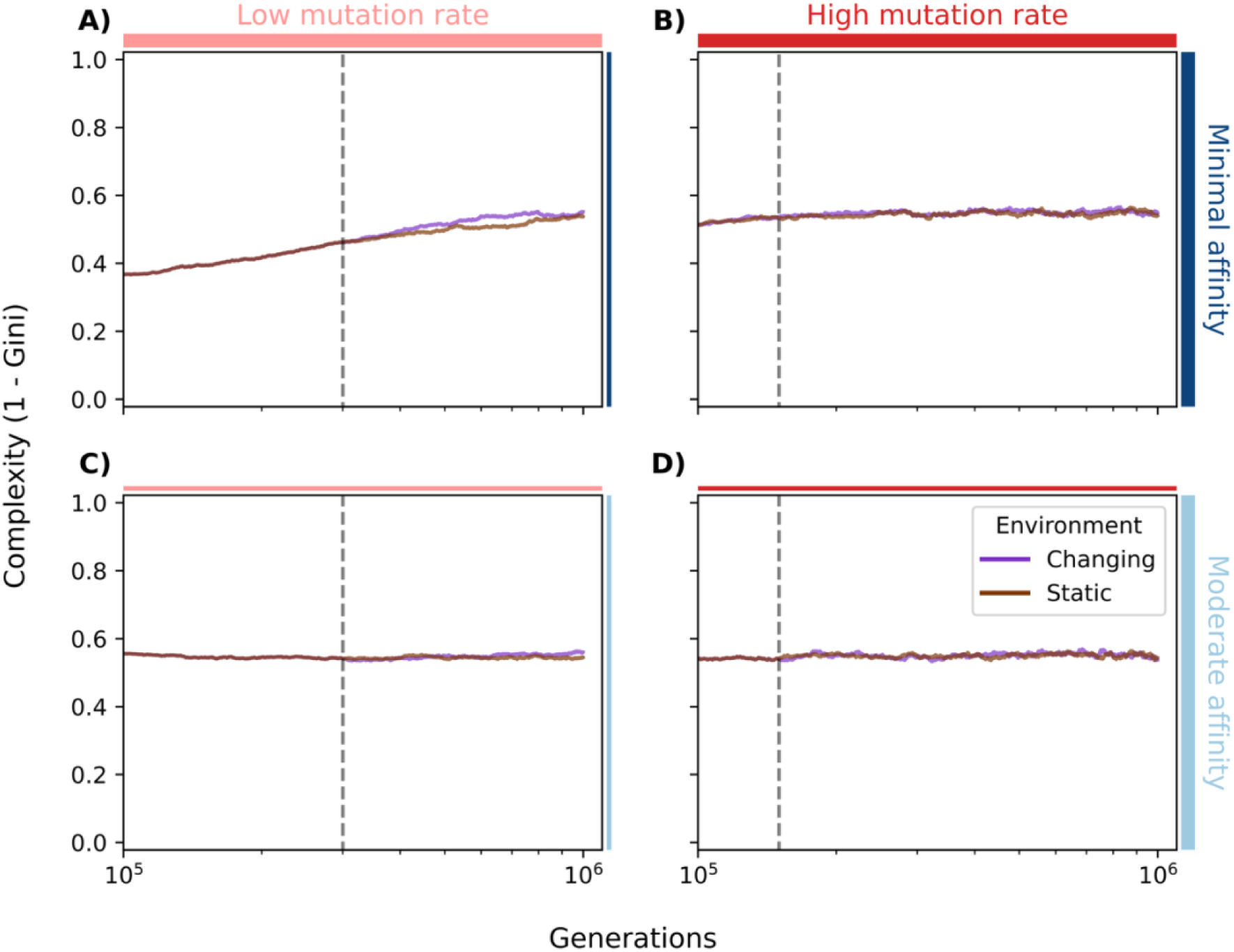
Non-recombining populations: changing environments accelerate complexity convergence unless plateau is already reached. Complexity (y-axis) shown as mean ± SEM (n = 10 replicates) up to 1 million generations (x-axis) under static (green) or changing (purple) environments. Panels show: **(A, C)** Low mutation rate, **(B, D)** high mutation rate, **(A, B)** minimal initial affinity, **(C, D)** moderate initial affinity.

**Supplementary Figure S8.**
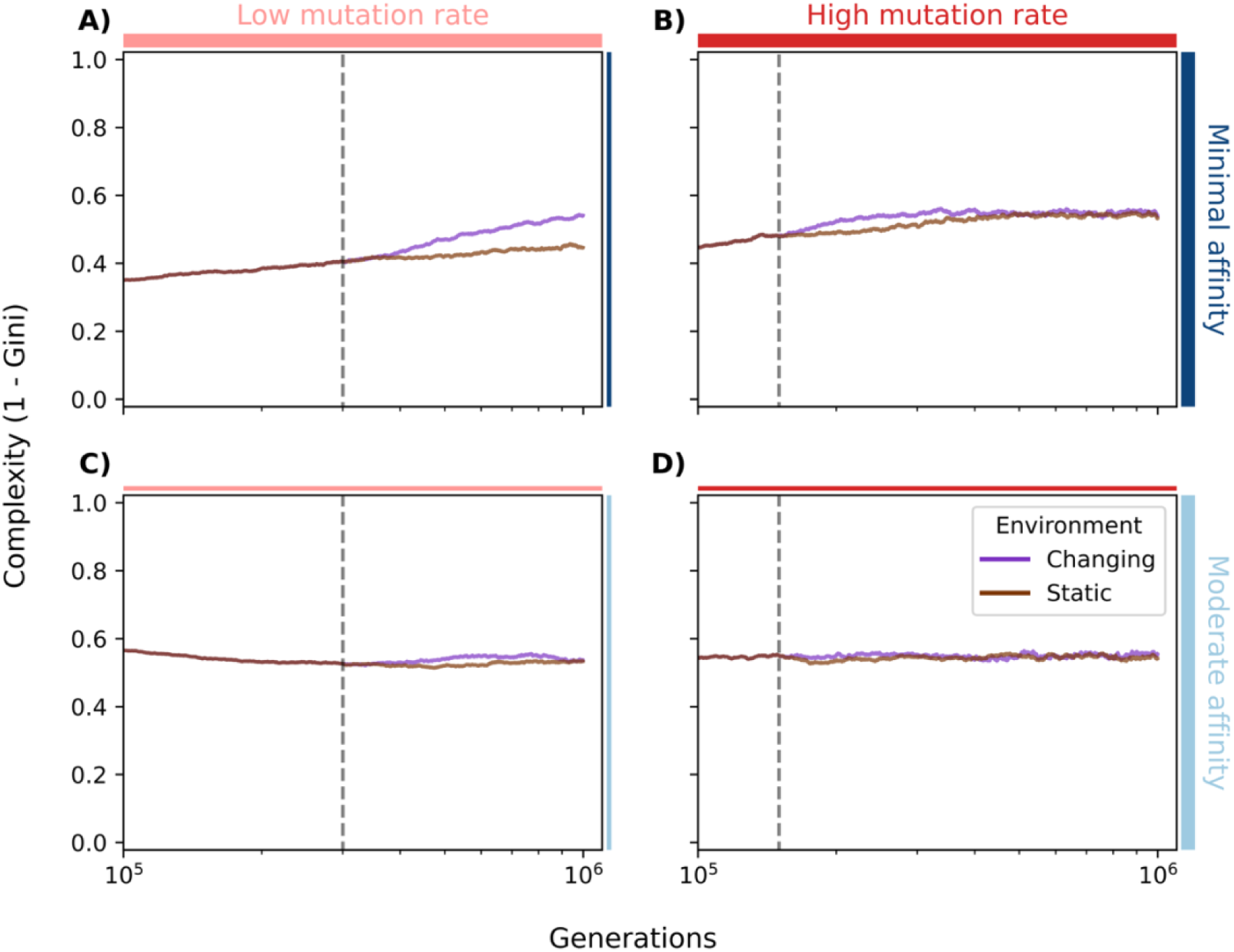
Recombining populations: changing environments accelerate complexity convergence unless plateau is already reached. Complexity (y-axis) shown as mean ± SEM (n = 10 replicates) up to 1 million generations (x-axis) under static (green) or changing (purple) environments. Panels show: **(A, C)** Low mutation rate, **(B, D)** high mutation rate, **(A, B)** minimal initial affinity, **(C, D)** moderate initial affinity.

**Supplementary Figure S9.**
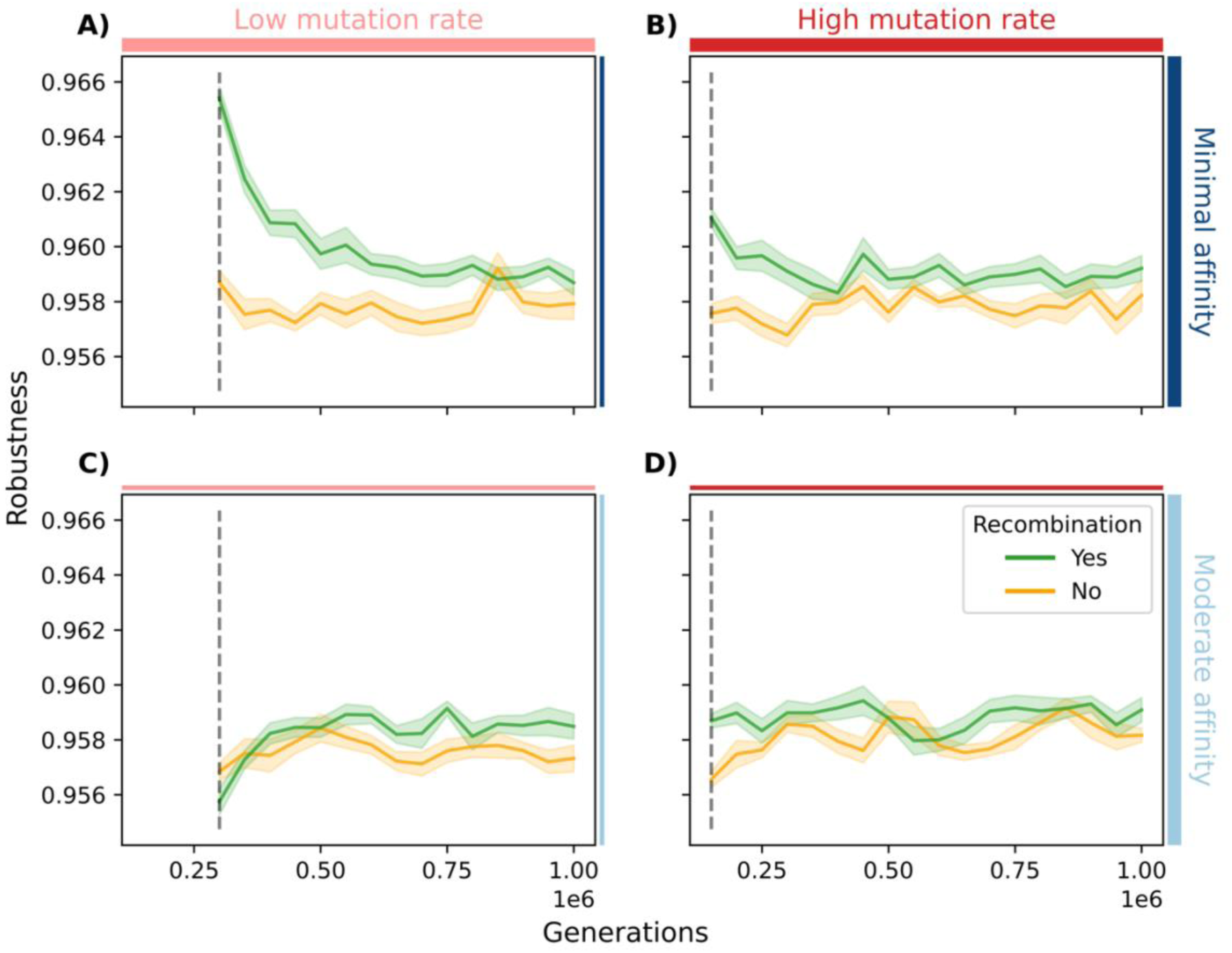
Recombining populations tend to evolve higher robustness than non-recombining populations in changing environments. Mutational robustness (y-axis) shown as mean ± SEM (n = 10 replicates) for 1 million generations (x-axis). Populations were evolved with (blue) or without (yellow) recombination. Panels show: **(A, C)** Low mutation rate, **(B, D)** high mutation rate, **(A, B)** minimal initial affinity, **(C, D)** moderate initial affinity. Environmental changes begin at the generations marked by vertical dashed lines.

**Supplementary Figure S10.**
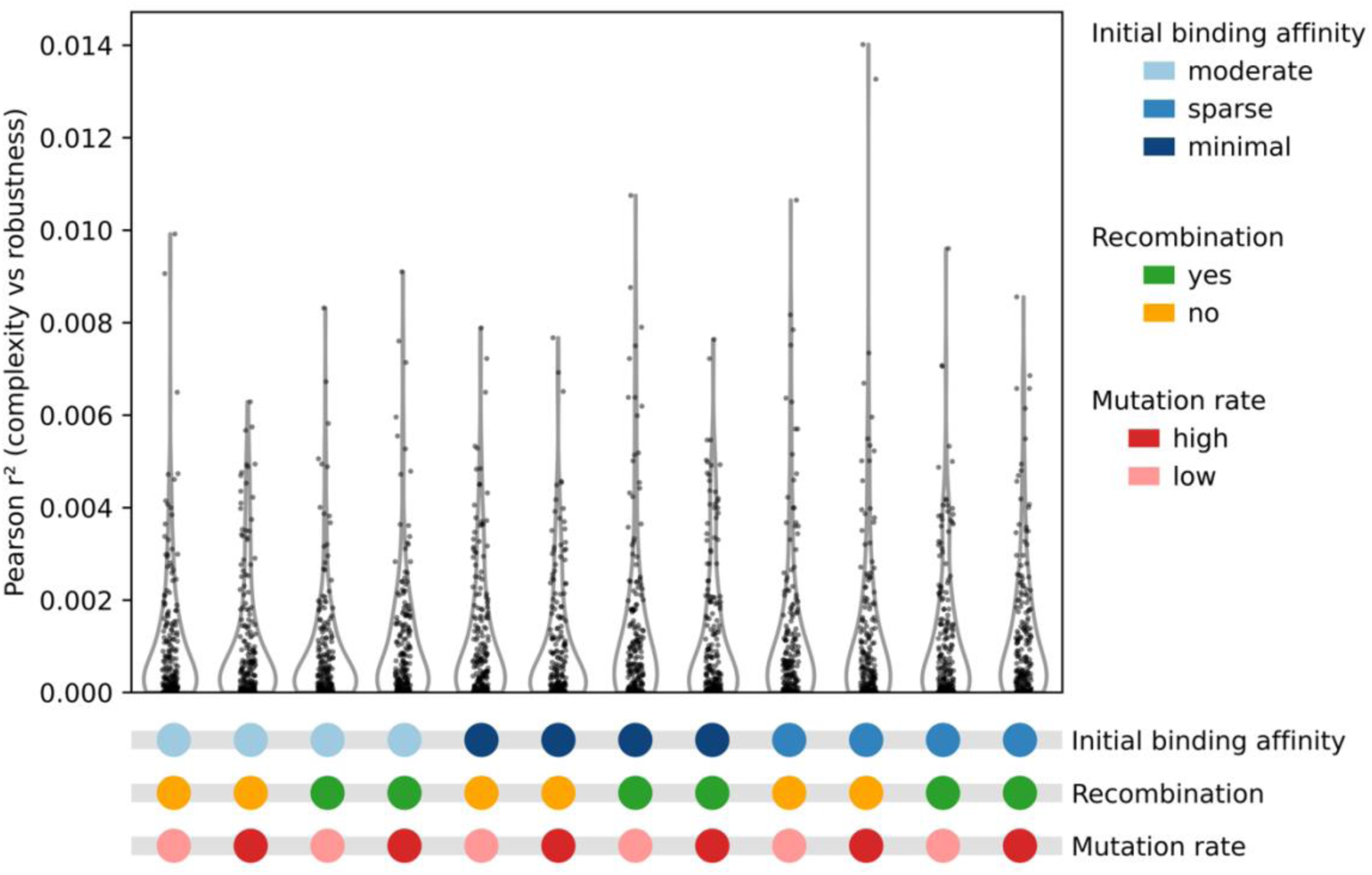
Complexity and robustness are uncorrelated. Pearson’s *r*^2^ values (y-axis) between complexity and robustness across all test conditions (x-axis), calculated individually for each population of 1,000 individuals (dots). Correlation was tested every 50,000 generations for each of 10 replicate populations per condition (n = 200 tests).

